# Model-based learning deficits in compulsivity are linked to faulty representations of task structure

**DOI:** 10.1101/2020.06.11.147447

**Authors:** Tricia X.F. Seow, Redmond O’Connell, Claire M. Gillan

## Abstract

Individuals with higher levels of compulsivity exhibit poorer performance on tasks that require model-based planning but the underlying causes have yet to be established. Here, we sought to determine whether these deficits stem from impoverished action-outcome relational knowledge (i.e. issues building an accurate model of the world) and/or an inability to translate models into action. 192 participants performed a two-step reinforcement learning task with concurrent EEG recordings. We found that representations of task-relevant action-outcome associations reflected in reaction time and parietal-occipital alpha-band power were stronger in individuals whose decisions were more model-based, and critically, were weaker in those high in compulsivity. At the time of choice, mid-frontal theta power, a general marker of cognitive control, was also negatively associated with compulsivity, but not model-based planning. These data suggest that model-based planning deficits in compulsive individuals may arise from failures in building an accurate model of the world.

## Introduction

Compulsive behaviour manifests as actions that are autonomous, out-of-control and repetitive, often leading to adverse and functionally impairing outcomes (Robbins, Gillan, Smith, de Wit, & Ersche, 2012). This symptomology is characteristic of psychiatric disorders like obsessive-compulsive disorder (OCD) and addiction, and is thought to arise from an imbalance between the two modes of action control (Gillan & Robbins, 2014): (i) goal-directed, model-based, behaviour that relies on knowledge of how actions lead to specific outcomes and (ii) rigid habits, that depend on more reflexive stimulus-response associations that form slowly over time (Balleine & O’Doherty, 2010; Dickinson, 1985). Up until now, the compulsivity literature has focused largely on testing the hypothesis that habitual behaviours dominate due to a dysfunctional imbalance in the competitive interactions between these two systems (Gruner, Anticevic, Lee, & Pittenger, 2016; Lee, Shimojo, & O’Doherty, 2014). In recent years however, evidence has begun to emerge that rather than being solely a failure in the arbitration between these two decision systems (i.e. both encouraging habit expression and discouraging planning), compulsivity may be driven primarily by impairments in goal-directed control. For example, imaging work has shown that behavioural insensitivity to rapid changes in outcome value (“outcome devaluation”) and habitual urges in OCD are associated with the hyperactivation of a brain area linked to goal-directed associative learning, the caudate nucleus, but not habit-related regions (Gillan et al., 2015). Moreover, OCD patients underperform on tasks that require prospective planning (Gillan, Morein-Zamir, Kaser, et al., 2014), and this is perhaps best exemplified by deficits in performance of the two-step reinforcement task (Voon et al., 2015). In this task, ‘model-based’ learning is operationalised as the extent to which individuals make decisions using knowledge of how their actions relate to subsequent events (Daw, Gershman, Seymour, Dayan, & Dolan, 2011; Daw, Niv, & Dayan, 2005). Not only are OCD patients impaired at this task, so too are individuals diagnosed with other compulsive disorders such as binge-eating disorder and addiction (Voon et al., 2015). Beyond case-control comparisons, more recent work has shown that this dysfunction is best captured by a continuum of compulsivity that is evident in both general population samples (Gillan, Kosinski, Whelan, Phelps, & Daw, 2016) and in diagnosed patients with or without a compulsive diagnosis (Gillan et al., 2019).

Despite these advances in our understanding of the nature of the deficits experienced by compulsive individuals, it remains unclear what drives model-based planning problems in compulsivity. Unarguably a multifaceted cognitive capacity, model-based planning depends upon several concurrent functions that include: (i) the construction and maintenance of an internal model (i.e. a representation of the environment, and more specifically, knowledge of relevant action-outcome relationships and state-state transitions), which is a pre-requisite for (ii) the implementation of this model in behaviour through prospective planning. Model-based failures could theoretically stem from problems in mechanisms underlying either component (in addition to several others that are not the focus of the present paper). Though direct tests to resolve this have been lacking, it has been suggested that OCD patients might have issues solely with implementation. For example, one study showed that patients have deficits in goal-directed decision making even when they have the requisite explicit knowledge of simple action-outcome contingencies (Gillan, Morein-Zamir, Urcelay, et al., 2014). However, other studies have suggested this might be a ceiling effect, revealing problems in learning action-outcome associations in both OCD and addiction using paradigms that require subjects to learn more numerous and/or taxing contingency structures (Ersche et al., 2016; Gillan et al., 2011). Moreover, this lack of action-outcome contingency knowledge was shown to correlate with failures in goal-directed control in OCD (Gillan et al., 2011). Overall, the evidence from these studies remains equivocal, because these tasks were (i) not designed to assess subjects’ ability to represent the task environment and (ii) used devaluation-style tasks that conflate deficits in goal-directed control with increases in stimulus-response habit learning.

In an adjacent literature, studies of metacognition have begun to investigate if internal representations of cause and effect may be compromised in compulsivity. For example, individuals high in compulsivity exhibit over-confidence in perceptual decision making tasks and may also have some difficulty in knowing when they are right versus wrong (Rouault, Seow, Gillan, & Fleming, 2018). A subsequent investigation of metacognition with respect to reinforcement learning also found that compulsive individuals were over-confident, but also that their confidence did not update appropriately in response to new evidence and did not inform their behaviour as strongly as those low in compulsivity (Seow & Gillan, 2020). Although not studied in the context of model-based planning specifically, these metacognitive findings suggest the possibility that in compulsivity, model-based planning failures may stem from fundamental issues in acquiring and maintaining an accurate internal model of the environment. Several studies have begun to probe healthy subjects’ ability to represent features of the task environment (i.e. action-outcome associations) and how they relate to model-based performance. For instance, one study showed that subjects who evoked the strongest neural representation of a future state also showed greater model-based planning (Doll, Duncan, Simon, Shohamy, & Daw, 2015). Others have shown that after making an initial choice, subjects’ reaction times (Shahar et al., 2019) and the way they move the computer mouse (Konovalov & Krajbich, 2020) can both be used to capture their knowledge of task structure. To date, no study has examined neural or behavioural representations of task structure in compulsive individuals as they perform a model-based planning task.

The present study aimed to fill this gap—testing if compulsivity is characterised by a disruption in constructing and maintaining an accurate representation of the task environment, or more simply the use/implementation of this ‘model’ in their choices. To do this, we used electroencephalography (EEG) to concurrently track neural signatures of state transition knowledge (P300 and posterior alpha) and of cognitive control (mid-frontal theta) as 192 subjects performed a two-step reinforcement learning task (***Figure 1***) (Daw et al., 2011, 2005). With single-trial regression analyses, we sought to characterise reaction time and several candidate neural correlates of the representation and implementation of the mental model linked to individual differences in model-based planning, and test if these same signatures were compromised in those high in compulsivity.

**Figure 1.**
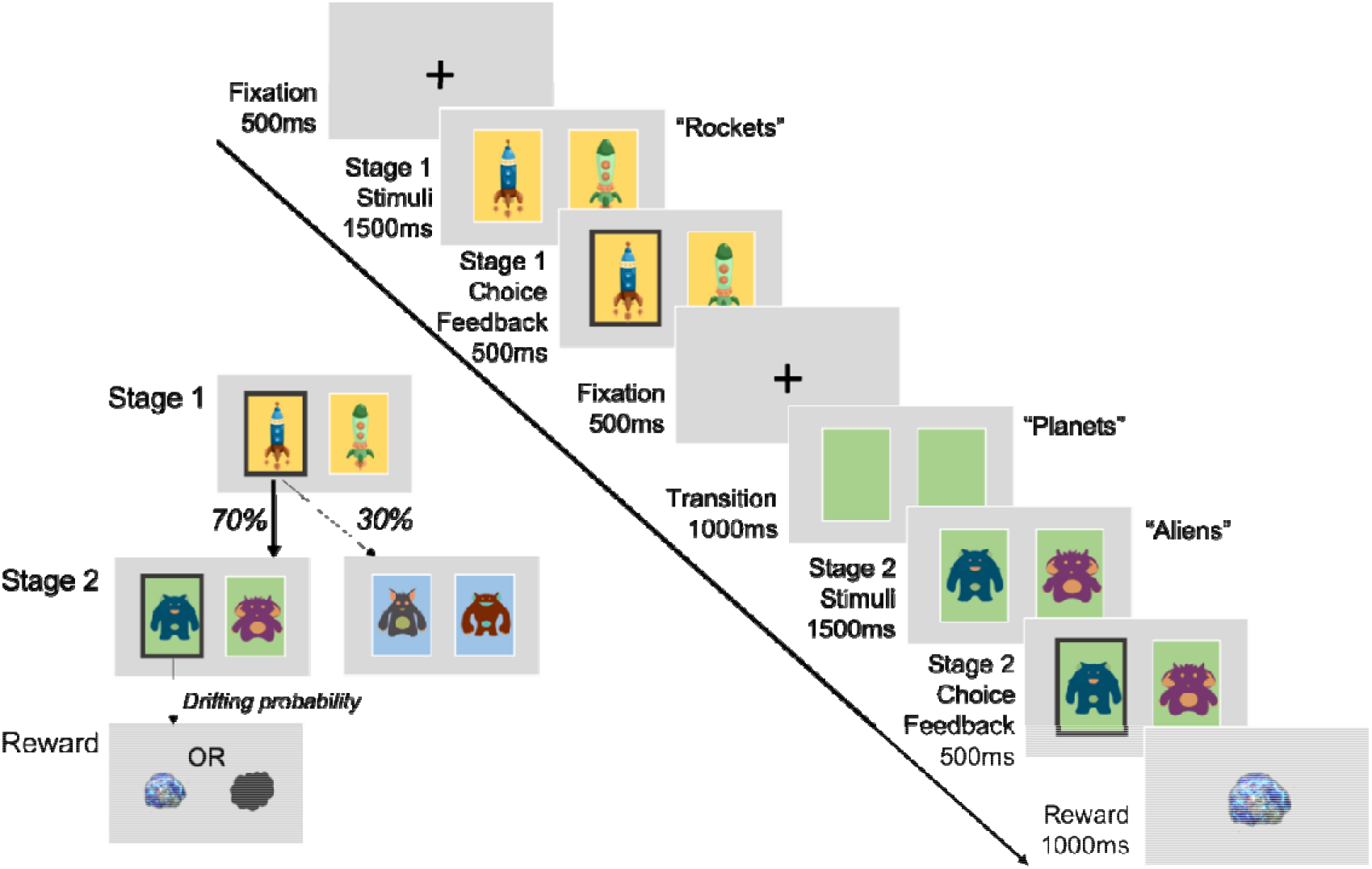
Two-step reinforcement learning task. Paradigm consists of two stages where participants take a rocket that has a common (70%) or rare (30%) transition to one of two second stage planets (states). Aliens on these planets each have a unique probability of reward (‘space treasure’ (reward) or ‘space dust’ (non-reward)) that drifts slowly throughout the entire experiment. Participants have to take into consideration the task transition structure and their history of rewards to make choices that maximise reward. The sequence of events as presented for EEG is the same as that of Eppinger et al. (2017), except they included a manipulation of transition probabilities in their study (comparing 60/40% to 80/20%) and used a longer choice window (2000ms).

## Results

### Compulsivity and model-based planning

Regression analysis of choice behaviour on the two-step task revealed a significant interaction between Reward and Transition (*β* = 0.20, *standard error (SE)* = 0.03, *p* < 0.001), indicating clear evidence for model-based planning in this sample. Individual subject coefficients for this interaction term were extracted and used as an individual difference measure for EEG analysis (split half-reliability was *r* = 0.71). Consistent with prior work, there was also evidence for model-free learning, where subjects were more likely to repeat choices if they were followed by reward (main effect of Reward: *β* = 0.55, *SE* = 0.05, *p* < 0.001), and an overall tendency to repeat choices from one trial to the next (Intercept: *β* = 1.46, *SE* = 0.07, *p* < 0.001). Importantly, we replicated prior work in finding that individual differences in compulsivity and intrusive thought (hereafter: ‘compulsivity’) were associated with reduced model-based planning (*β* = −0.07, *SE* = 0.04, *p* = 0.05) (***Figure 2A***), while anxious-depression (*β* = 0.05, *SE* = 0.04, *p* = 0.14) and social withdrawal were not (*β* = −0.01, *SE* = 0.04, *p* = 0.73).

**Figure 2.**
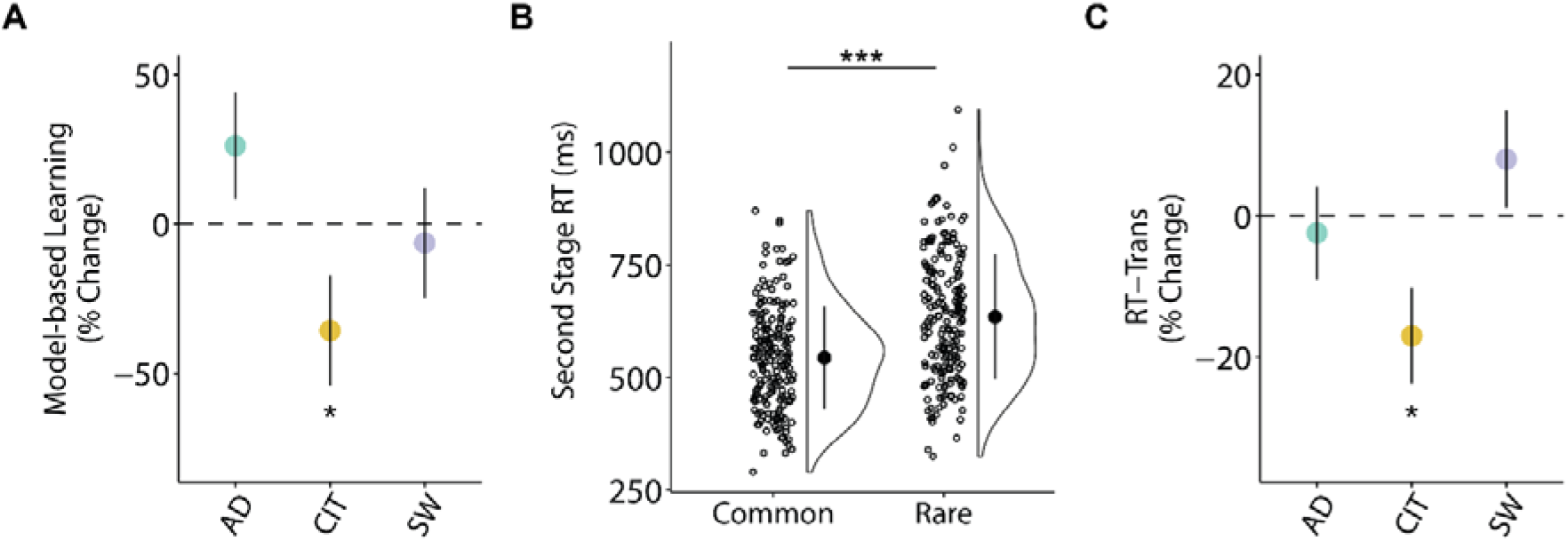
Model-based behaviour and reaction times in compulsivity. (**A**) Model-based control estimated by a logistic regression of choice behaviour with one-trial back reward and transition. Regressions were conducted in a model with all three dimensions: ‘anxious-depression’ (AD), ‘compulsivity and intrusive thought’ (CIT) and ‘social withdrawal’ (SW). Model-based control is reduced high compulsive individuals. (**B**) Participants have on average a longer mean response time (RT) at second stage choice after a rare transition than a common one (t_191_ = 16.16, 95% Confidence Interval (CI) [79.85 102.05], p < 0.001). Circles in raincloud plot (Allen, Poggiali, Whitaker, Marshall, & Kievit, 2019) depict mean RT of rare or common trials for each individual, with black marker indicating grand average mean and standard deviation (SD). (**C**) RT difference between transition type (RT-Trans) is diminished in high compulsive individuals. For (**A**) & (**C**), error bars denote standard error. The Y-axes indicate the percentage change in model-based planning/RT-Trans as a function of 1 SD of psychiatric dimension scores. *p ≤ 0.05, ***p < 0.001.

### Reaction time (RT) sensitivity to task structure

Someone who is aware of the task structure should expect to be presented with the second stage state that is most commonly associated with their first stage choice. As such, when a rare transition occurs, they will require more time to respond and ‘re-plan’ (Shahar et al., 2019). In line with this previous work, we hypothesised that participants would have a slower RT after a rare versus common transition and that this difference would be greater in participants who exhibit the most model-based behaviour. We found support for both hypotheses; participants had a slower mean RT for rare versus common trials after transition (*β* = 47.15, *SE* = 2.85, *p* < 0.001) (***Figure 2B***) and this effect was larger in those with higher levels of model-based control (*β* = 7.36, SE = 1.83, *p* < 0.001). Crucially, we found that this effect was reduced in high compulsive individuals (*β* = −8.03, *SE* = 3.19, *p* = 0.01) (***Figure 2C***). Prior studies using this task did not test for an association between compulsivity and this RT cost, but the data is readily available. To test the robustness of this finding, we therefore re-analysed a prior dataset (N = 1413) collected entirely online (Gillan et al., 2016) using a similar variant of the two-step task and the same measure of compulsivity. We replicated this effect (*β* = −5.80, *SE* = 1.58, *p* < 0.001). This is, to our knowledge, the first evidence that compulsivity is associated with a deficit in distinguishing rare from common transitions, a fundamental constituent of the mental-model required to engage in model-based planning.

### P300 sensitivity to task structure

The P300 or P3b has well-established sensitivity to stimulus probability, exhibiting larger peak amplitudes for less probable stimuli (Polich & Margala, 1997). Prior research in healthy humans thus hypothesised that the P300 may be a marker of sensitivity to state transitions on the two-step task, though these studies have yielded inconsistent results, with some finding greater P300 amplitudes for rare versus common transitions (Sambrook, Hardwick, Wills, & Goslin, 2018; Shahnazian, Ribas-Fernandes, & Holroyd, 2019) and one finding the opposite (Eppinger, Walter, & Li, 2017). Here, we examined the second stage stimulus-locked P300 and found a significant main effect of transition type (*β* = 0.15, *SE* = 0.07, *p* = 0.03), consistent with Sambrook et al. (2018) and Shahnazian et al. (2019) whereby greater P300 amplitude was observed after rare versus common transitions (***Figure 3***). However, this differential rare versus common signal was not larger in individuals high in model-based planning (*β* = 0.07, *SE* = 0.08, *p* = 0.35), nor did it show any association to compulsivity (*β* = 0.09, *SE* = 0.08, *p* = 0.24).

**Figure 3.**
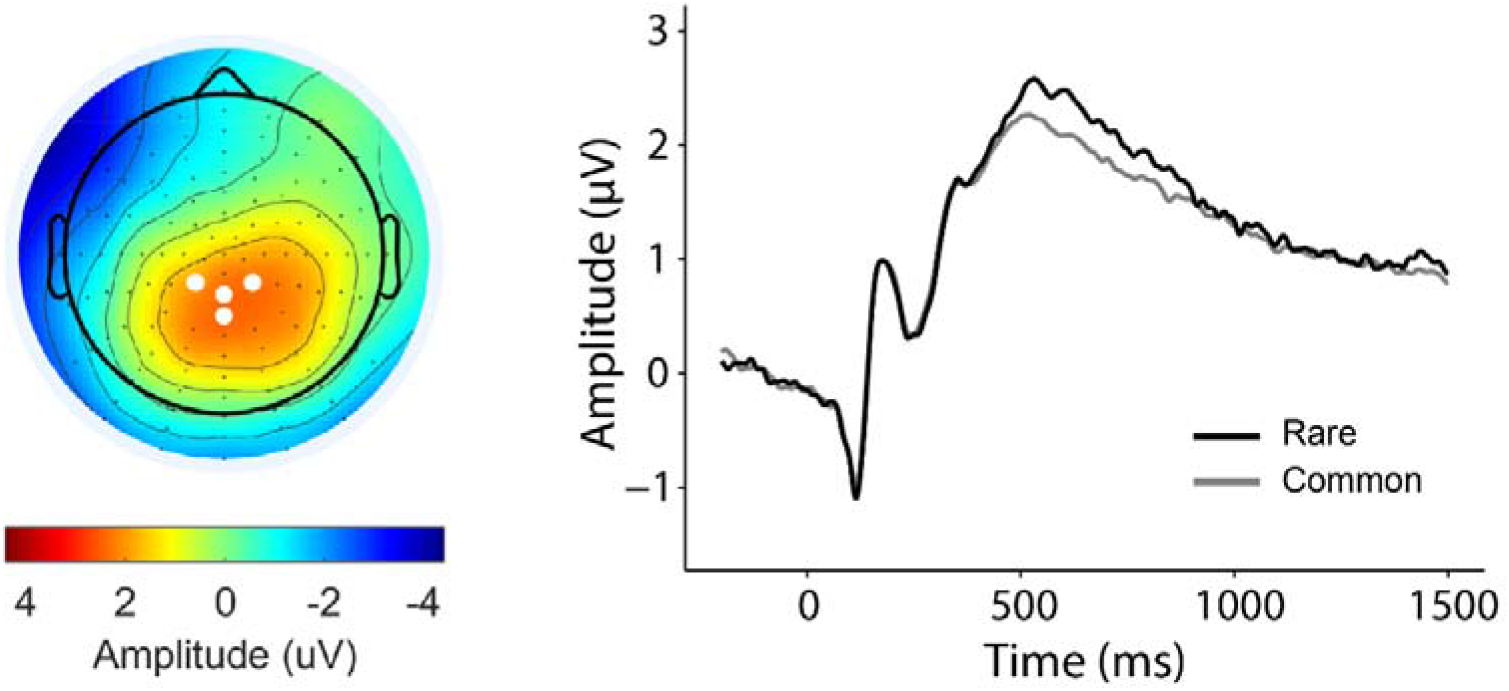
Second stage stimulus-locked P300 and transition type. Grand average waveforms of rare and common trials locked to second stage stimuli (aliens). Waveform is baselined −200ms to 0ms. The mean amplitude for stimulus-locked P300 was obtained over 4 centro-parietal electrodes (D16 (CP1), A3 (CPz), B2 (CP2), A4) as indicated by the white dots in the topography plot.

Recently, it has been demonstrated that the P300 component traces the accumulation of evidence for decision making and, as such, is time-locked to choice commitment (O’Connell, Dockree, & Kelly, 2012; Twomey, Murphy, Kelly, & O’Connell, 2015). This raises the possibility that the stimulus-locked signal measurements favoured in previous studies of the two-step task may have yielded cross-condition effects that were partly or entirely determined by RT differences. In light of these considerations, we complemented the stimulus-locked analyses with an examination of response-locked signal measurements. When we repeated the analysis using response-locked P300 amplitude, we found that the transition effect was no longer significant and its direction was in fact reversed (*β* = −0.09, *SE* = 0.08, *p* = 0.23) (***Supplemental Figure S3***). Again, there was no association with model-based planning (*β* = −0.05, *SE* = 0.08, *p* = 0.49) or compulsivity (*β* = 0.04, *SE* = 0.09, *p* = 0.67). We also examined the build-up rate of the response-locked P300 as a measure of how quickly evidence for the decision was accumulated (Kelly & O’Connell, 2013). The build-up rate was steeper for common versus rare trials (*β* = −0.001, *SE* = 0.0004, *p* = 0.002) but this measure was again not linked to model-based planning (*β* = −0.0002, *SE* = 0.0004, *p* = 0.46) nor compulsivity (*β* = 0.0005, *SE* = 0.0004, *p* = 0.25). Thus, we concluded that the P300 may not provide the most reliable or sensitive measure of neural sensitivity to task structure.

### Alpha power sensitivity to task structure

As ERPs principally reflect activity changes that are short-lived and strictly time-locked to particular events (Makeig & Onton, 2012), we investigated if time-frequency measures such as alpha power (9-13Hz) would have a role in orchestrating more distributed neural computations of the sort required in model-based planning, and provide a superior and more sustained representation neural index of the task’s transition structure. As general marker of mental activity and attention (Klimesch, 2012; Laufs et al., 2003), prior studies have found evidence for alterations in alpha power in OCD patients (Perera, Bailey, Herring, & Fitzgerald, 2019). Here, we reasoned that alpha power might be sensitive to transition structure given the increased mental activity required to re-plan when one arrives at an unexpected state.

Specifically, we examined if parietal-occipital alpha power locked to the second stage stimulus was able to distinguish between rare and common transitions across a series of time bins in our task. This allowed us to ascertain not just if participants showed sensitivity to task structure following a transition, but for how long they sustained that representation (e.g. as the made subsequent choices and received a reward). In line with our hypothesis, alpha power overall differentiated between the two transition types (*β* = 0.05, *SE* = 0.01, *p* < 0.001), such that parietal-occipital alpha was more suppressed after rare versus common transitions (***Figure 4A***). We found that over three rolling time bins beginning from the state transition (planet) (0ms to 1000ms: *β* = 0.04, *SE* = 0.02, *p* = 0.03) to the end of choice feedback (1000ms to 2000ms: *β* = 0.03, *SE* = 0.01, *p* = 0.03; 2000ms to 3000ms: *β* = 0.03, *SE* = 0.02, *p* < 0.05), individuals high in model-based control showed the largest alpha power differentiation (***Figure 4B***). Importantly, this same signature was negatively related to compulsivity, with a significant association observed at the time after state transition (0ms to 1000ms: *β* = −0.06, *SE* = 0.02, *p* = 0.007) (***Figure 4C***). Overall second stage alpha power was also associated with compulsivity (*β* = −0.20, *SE* = 0.05, *p* < 0.001), however, this effect was not related to model-based control (*β* = 0.04, *SE* = 0.05, *p* = 0.42) nor RT differences in transition types (*β* = −0.06, *SE* = 0.05, *p* = 0.20)—highlighting that it is the sensitivity of alpha to task structure, not alpha overall, that best tracks model-based performance at this task.

**Figure 4.**
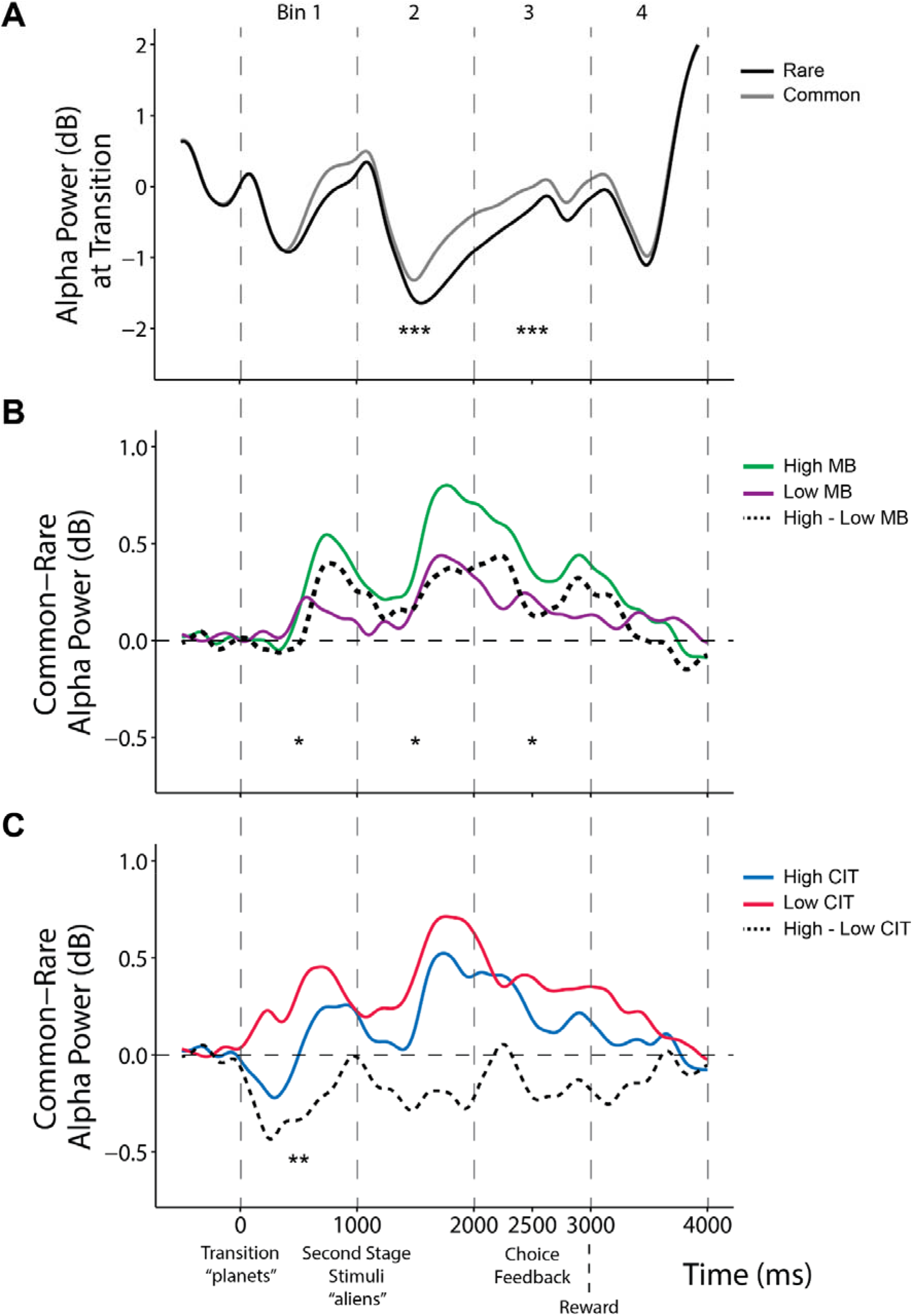
Stimulus-locked alpha power at transition. Alpha power was measured across 4 time bins of 1000ms each separated by vertical dashed lines, starting from the transition (0ms) until after reward (4000ms). (**A**) Grand average second stage alpha power waveforms between rare and common transitions. Continuous analyses revealed that alpha difference (rare – common) is significant in time bin 2-3 (all β > 0.06, SE < 0.02, p < 0.001). (**B**) Alpha power difference between transitions (common minus rare) is depicted above by comparing top/bottom 50^th^ percentile (N = 96 per group) of participants grouped by model-based estimates (MB). Continuous analyses revealed that alpha difference (rare – common) is enhanced for more model-based participants in time bins 1-3 (all β > 0.03, SE < 0.02, p < 0.05). (**C**) Alpha power difference between transitions (common minus rare) comparing top/bottom 50^th^ percentile (N = 96 per group) of participants grouped by or compulsivity (CIT). Continuous analyses revealed that alpha difference (rare – common) is diminished for more compulsive participants in time bin 1 (β = −0.06, SE = 0.02, p = 0.007). Stars in time bins indicate significance from continuous analyses. *p < 0.05, **p < 0.01, **p < 0.001.

Notably, control analyses also indicate that the alpha transition effect is not explained by RT confounds (***Supplemental Figure S5***) and second stage theta-power (which we examine later in the context of cognitive control at first stage) did not exhibit the same results (i.e. transition effect is specific to alpha-band activity) (***Supplemental Figure S6***). Finally, when we examined the association between alpha power sensitivity to transition structure and the entire set of nine psychiatric questionnaire scores, we found that the effect was present in more than one measure: diminished in those with elevated OCD (*β* = −0.03, *SE* = 0.01, *p* = 0.006) and eating disorder symptoms (*β* = −0.03, *SE* = 0.01, *p* = 0.05) (***Figure 5***). The transdiagnostic analysis thus improved the specificity of the association—showing this deficit tracked compulsivity and not anxious-depression or social withdrawal.

**Figure 5.**
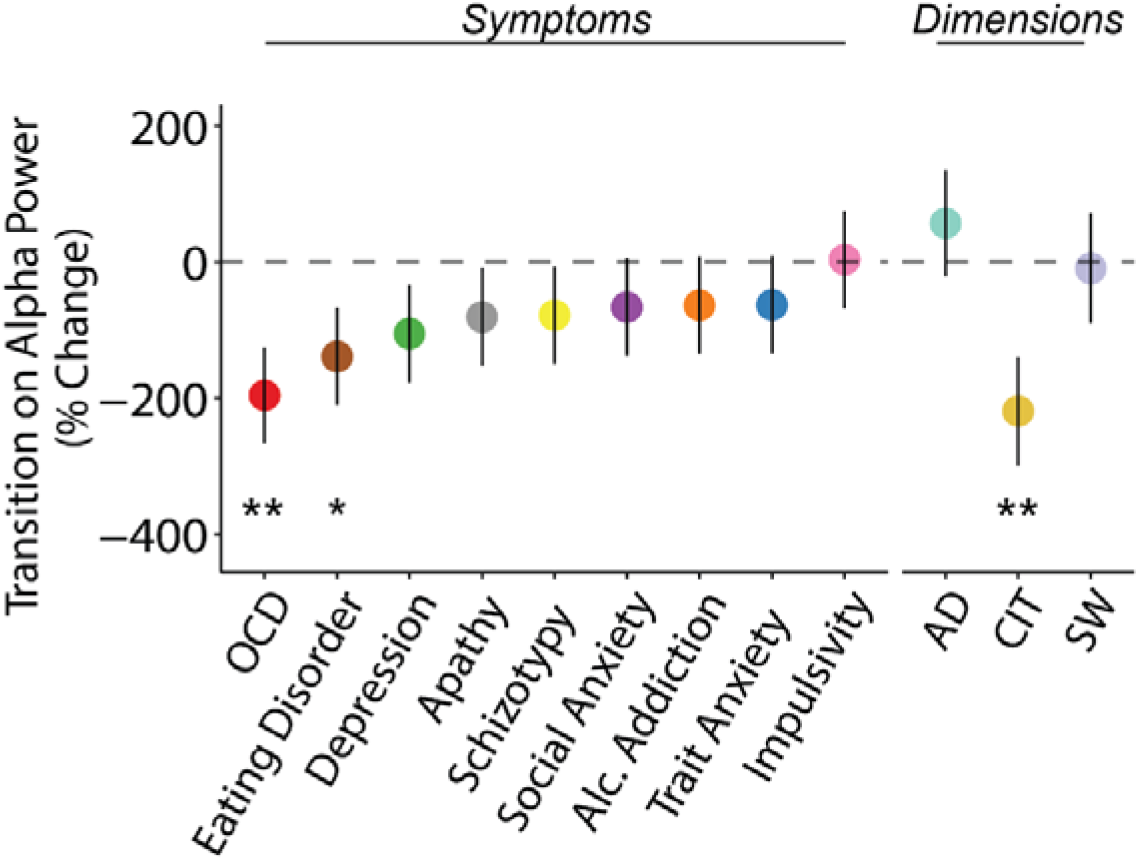
Second stage alpha power sensitivity to transition at time bin 1 (0ms to 1000ms) with psychiatric symptoms and dimensions (AD: ‘anxious-depression’, CIT: ‘compulsive behaviour and intrusive thought’, SW: ‘social withdrawal’). Alpha power differentiating rare versus common transitions was associated with both OCD and eating disorder symptoms. The transdiagnostic analysis showed the effect was captured by a compulsive dimension (CIT). The Y-axis show the percentage change in alpha power sensitivity to transition type (%) as a function of 1 SD increase of psychiatric questionnaire/dimension scores. Error bars denote standard errors. *p ≤ 0.05, **p < 0.01.

### Theta power at the time of choice

Finally, moving beyond participants’ awareness of the transition structure of the task, we tested for a more general disruption in cognitive control, assayed as theta (4-8Hz) power during the first stage choice—the crucial time when model-based planning manifests in behaviour. Mid-frontal theta power is a well-established EEG signature of exerting ‘cognitive control’ over lower level impulses (Cavanagh & Frank, 2014; Sauseng, Griesmayr, Freunberger, & Klimesch, 2010), including Pavlovian biases (Cavanagh, Eisenberg, Guitart-Masip, Huys, & Frank, 2013). We therefore considered theta power as a candidate signature associated with implementing model-based decisions and overriding more habitual, model-free choices, and as such hypothesized that theta would be positively associated with model-based planning and negatively linked to compulsivity.

Contrary to this, we found that theta power during choice was not significantly associated with model-based planning (*β* = 0.02, *SE* = 0.01, *p* = 0.11), though the trend was in the expected direction. Despite this, we found an overall effect of lower theta at the time of choice in individuals high in compulsivity (*β* = −0.03, *SE* = 0.01, *p* = 0.03) (***Figure 6A***). Similar to alpha power modulations, reduced theta power at the first stage was linked to more than one questionnaire score—schizotypy (*β* = −0.03, *SE* = 0.01, *p* = 0.01), depression (*β* = −0.03, *SE* = 0.01, *p* = 0.02) and OCD (*β* = −0.02, *SE* = 0.01, *p* = 0.03). Again, these effects were captured by the compulsive dimension (*β* = −0.03, *SE* = 0.01, *p* = 0.03) (***Figure 6B***). Theta power was associated with RT sensitivity to transition structure such that those subjects who had higher theta power during their first stage choice had larger differences in their RT between rare and common transitions at the second-stage (*β* = 0.03, *SE* = 0.01, *p* = 0.001).

**Figure 6.**
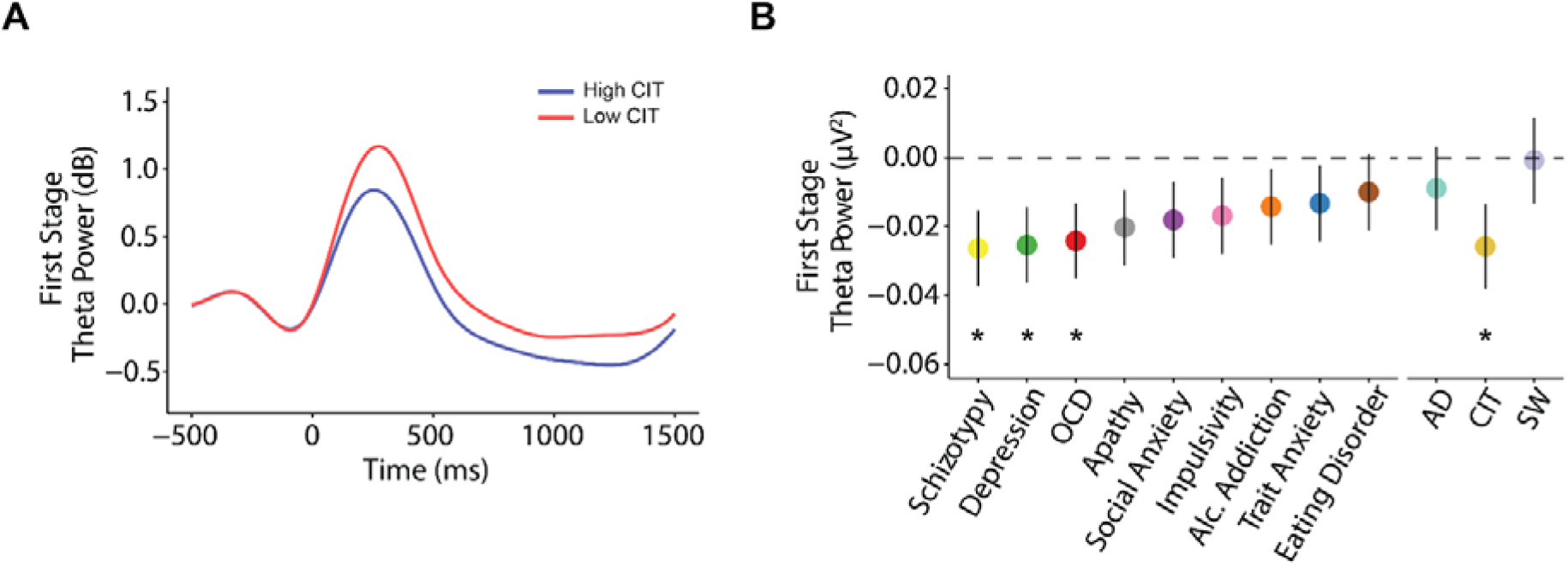
First stage theta power with psychiatric symptoms and dimensions (AD: ‘anxious-depression’, CIT: ‘compulsive behaviour and intrusive thought’, SW: ‘social withdrawal’). (**A**) Grand average waveforms of theta power comparing the top/bottom 50th percentile (N = 96 per group) individuals based on their model-based (MB)/compulsivity (CIT) estimates. Single trial analyses (with all participants) indicate high compulsive individuals exhibit a decrease in theta power (β = −0.03, SE = 0.01, p = 0.03). (**B**) Reduced theta power at first stage was linked to several questionnaire scores but the effect was ultimately specific to compulsivity. The Y-axis shows the change in theta power (μV^2^) as a function of 1 SD increase of psychiatric questionnaire/dimension scores. Error bars denote standard errors. *p < 0.05.

By way of control analysis, we tested if alpha power at first stage was associated with compulsivity (*β* = −0.56, *SE* = 0.03, *p* = 0.002), model-based planning (*β* = 0.12, *SE* = 0.02, *p* = 0.45) or RT differences in transition types (*β* = −0.02, *SE* = 0.16, *p* = 0.92), but none were significant (***Supplemental Figure S4***).

## Discussion

Model-based planning deficits linked to compulsivity have been theorised to arise from issues with the balance/arbitration between competing model-based and model-free influences during choice (Gillan & Robbins, 2014; Gruner et al., 2016; Lee et al., 2014; Lloyd & Dayan, 2019). We tested if these behavioural impairments might also stem from dysfunctions in generating an accurate model of the world (here, the representation of action-state transitions in the two-step task). We used EEG and reaction time analysis to investigate if model-based deficits in compulsivity reflect an impoverished internal model of the task, difficulties in executing that model, or both. We found that internal representations of cause and effect were diminished in compulsivity—high compulsive individuals lacked sensitivity to state transition probabilities, evidenced in their reaction times (RT) and parietal-occipital alpha power suppression. We also observed that mid-frontal theta, a general marker of cognitive control, was reduced when subjects made their initial choices on this task, though this signal bore a less clear relation to model-based behaviour than the alpha findings. These findings have important implications for refining theories of compulsivity and future research examining the potential role of metacognitive processes in abnormalities in model-based planning.

An analysis of RT provided the first evidence that high compulsive individuals were less aware of state transitions. In line with prior research, the subject group as a whole exhibited longer RTs following rare transitions (Shahar et al., 2019), which is presumed to reflect the fact that one needs to adjust to this unlikely event and ‘re-plan’ their next choice. In line with this account, we found that individuals who had higher levels of model-based planning performance showed the largest RT differences following rare versus common transitions. Crucially, the opposite was true of compulsivity, with the most compulsive individuals showing the smallest difference in RT for these trial types. Notably, this finding was robust—we replicated the same effect in a former dataset (with N = 1413) tested online (Gillan et al., 2016). Thus, compulsivity appears to be associated with having diminished awareness of the transition structure of the task, a key requirement for engaging in model-based planning on this task.

Moving beyond behaviour, analysis of alpha power following these state transitions revealed a strikingly similar picture. Much like RT, alpha desynchronization at the second stage of this task was sensitive to whether a rare or common transition had occurred. Specifically, rare transitions were associated with greater alpha desynchronization compared to common trials, possibly reflecting the greater mental effort required on these trials to call to mind the action values associated with options the individual was not expecting to see. Consistent with this interpretation, previous studies using n-back paradigms have shown parieto-occipital alpha is more suppressed when working memory load increases (Pesonen, Hämäläinen, & Krause, 2007; Stipacek, Grabner, Neuper, Fink, & Neubauer, 2003). Individual difference analysis demonstrated that this difference in alpha desynchronization had important behavioural correlates, where those individuals who were highest in model-based planning showed the largest differences in alpha power for rare versus common transitions. Importantly, the effect for compulsivity was reversed—higher levels of compulsivity were associated with less of a distinction in alpha power for rare versus common transitions. Building upon the reaction time data, this is the first *neural* evidence that compulsivity may be characterised by failures in representing the kind of causal action-state relations necessary to behave in a model-based manner.

Our data do not exclude the possibility that compulsive individuals also face issues with implementing model-based planning in situations when they might have the requisite state knowledge. Mid-frontal theta reflects a common mechanism for executing adaptive control in a variety of contexts (Cavanagh, Zambrano-Vazquez, & Allen, 2012) and more specifically for selecting between competing options, including the suppression of distracting stimuli when focused attention is required (Nigbur, Ivanova, & Stürmer, 2011). As theta was a good candidate neural signature of model-based deliberation, we thus tested if there was evidence for reduced theta power in compulsive individuals at the crucial time of first stage choice, when model-based and model-free action values putatively compete for control. We found that theta power was indeed reduced in high compulsive individuals during first stage choice, though evidence of a link between increased theta and increased model-based planning itself was not compelling in the present study. However, it is interesting to note that greater theta was associated with greater RT differentiation of transition types, which could suggest that theta power at the time of choice in part reflects the simulation of future states, a form of implementation that would naturally be elevated in those with the strongest mental model. A previous study that examined task-related theta power in OCD found that patients exhibited lower theta power during tasks that require inhibitory regulation (Min, Kim, Park, & Park, 2011), suggesting that reduced levels observed here in high compulsive individuals might reflect a failure to inhibit competing model-free action values. It is equally possible, however, that reduced theta might reflect a lack of the existence of competing signals. For instance, if individuals fail to represent a model of the task environment, then one might argue that the need to engage cognitive control is necessarily reduced. Studies with Flanker tasks suggest that theta is linked to conflicts in both sensory perception and response control (Nigbur, Cohen, Ridderinkhof, & Stürmer, 2012); future work may examine these processes in compulsivity.

Previous EEG studies of the two-step task (Eppinger et al., 2017; Sambrook et al., 2018; Shahnazian et al., 2019) showed that the P300 was associated with state transitions. However, the inconsistent direction on the effects raises doubt as to how these differences should be interpreted. Recent literature conceptualises the P300 as an evidence accumulation process that builds towards a peak at choice time (Twomey et al., 2015) and as such variances in RT will influence the latency of the stimulus-locked P300 amplitude peak (Kelly & O’Connell, 2015). Our results comparing stimulus-locked and response-locked analysis approaches suggest that it is the build-up rate of the P300 that is sensitive to stimulus transitions and that previously reported modulations of its stimulus-locked amplitude are attributable to RT variances. Here however, we found that none of the analysed P300 metrics were predictive of individual differences in model-based planning. Shahnazian and colleagues also investigated theta (but not alpha) frequencies as potential neural correlates of state transitions related to model-based control on the two-step task (Shahnazian et al., 2019). We replicated their null results for theta here (***Supplemental Figure S6***).

In this study, we utilised a transdiagnostic compulsive dimension which was previously shown to provide the best mapping to model-based deficits in a general population sample tested online (Gillan et al., 2016). Here in our sample tested in-lab, we found that the modulations of alpha and theta power were non-specific when examined across the individual DSM-defined questionnaires, but the dimensional approach revealed more specific associations: impairments in the representation of the mental model and cognitive control were solely linked to the compulsive dimension (and not anxious-depression or social withdrawal phenotypes). A limitation of the current study is that we cannot directly address the applicability of our findings to diagnosed patients, but a recent patient study has shown that model-based deficits were more strongly linked with a dimension of compulsivity than OCD diagnosis status (Gillan et al., 2019). As such, the specific associations between dysfunctional cognition and compulsivity we delineate are likely clinically relevant.

Overall, our findings suggest that model-based difficulties in compulsivity are linked to an impoverished mental model of environmental contingency, assayed in both neural signals and reaction times. These findings may have implications for understanding how compulsive behaviours and obsessive beliefs develop in concert in disorders like obsessive-compulsive disorder. Clinical cognitive models of OCD have long presumed that compulsions are performed to reduce anxiety induced by obsessive beliefs (Matthews & Wells, 2008; Salkovskis & McGuire, 2003), in contrast to a more recent hypothesis suggesting that obsessions are post-hoc rationalisations to explain the performance of compulsive behaviour (Gillan & Sahakian, 2015). These data may suggest that the hard distinction between obsessions and compulsions might be less clear than these models propose. Failures in accurately representing the relationship between actions and their consequences, both implicitly and explicitly, may be a common source of both compulsive habitual behaviours in OCD and also faulty metacognitive beliefs that form the basis of obsessions. One might imagine that the less stable the representation of the world model is, the more likely a patient may develop faulty beliefs and rely on habitual representations.

## Materials and Methods

### Power estimation

We determined a minimum sample size from a prior study that investigated the association of goal-directed control (on a different task) with OCI-R scores from non-clinical participants who were also tested in-person (*r* = −0.26, *p* < 0.05) (Snorrason, Lee, de Wit, & Woods, 2016). The effect size indicated that N = 150 participants were required to achieve 90% power at 0.05 significance. Our final sample was larger than this to achieve the required power for another study that the same subjects participated in (Seow et al., 2019).

### Participants

N = 234 participants were tested, of whom 138 were female (58.97%) with ages ranging from 18 to 65 (mean = 31.42, standard deviation (SD) = 11.48) years. Majority of the participants were from the general public, recruited via flyers and online advertisements, while a tiny subset (N = 8) were attending group therapy for anxiety at St. Patrick’s University Hospital in Dublin, Ireland, included to enrich our sample in self-report mental health symptoms. All participants were ≥18 years (with an age limit of 65 years) and had no personal/familial history of epilepsy, no personal history of neurological illness/head trauma nor personal history of unexplained fainting. Subjects were paid €20 Euro (€10/hr) upon completion of the study. All study procedures were approved by Trinity College Dublin School of Psychology Research Ethics Committee.

### Procedure

Before presenting to the lab for in-person EEG testing, participants completed a brief at-home assessment via the Internet. They provided informed electronic consent, and submitted basic demographic data (age, gender), listed any medication they were taking for a mental health issue and completed a set of 9 self-report psychiatric questionnaires (see **Self-report psychiatric questionnaires, transdiagnostic dimensions & IQ**). During the in-person EEG session, participants completed two tasks: the modified Eriksen flanker task (Eriksen & Eriksen, 1974) and the two-step reinforcement learning task (Daw et al., 2011, 2005). Data from the former task data are presented elsewhere (Seow et al., 2019), but note that we also reported the basic association with compulsivity and model-based planning in that paper, which served to contextualise a null result. Once participants had completed both tasks, they completed a short IQ evaluation before debriefing. A subset of the participants (N = 110, 47%) completed a short psychiatric interview (Mini International Neuropsychiatric interview English Version 7.0.0; M.I.N.I.) (Sheehan et al., 1998) before the experimental tasks to establish their diagnostic status. Further details are in ***Supplemental Table S1.***

### Participant exclusion criteria

Several *a priori* exclusion criteria were applied to ensure data quality. Participants were excluded if they failed any of the following on a rolling basis: Participants whose/who (i) EEG data were incomplete (N = 5) (i.e. recording was prematurely terminated before the completion of the task) or corrupted (N = 2), (ii) EEG data which contained excessive noise (i.e. >70% EEG epochs from the individual failing epoch exclusion criteria, see **EEG recording & pre-processing**) (N = 4), (iii) responded with the same key in stage one >90% (n > 135 trials) of the time (N = 10), (iv) probability of staying after common-rewarded trials was significantly worse than chance, measured as <5% probability of fitting a binomial distribution with 50% (chance) probability and the total number of common-rewarded trials experienced by each subject (N = 11), (v) missed more than 20% of trials (n > 30 trials) (N = 3), and (vi) incorrectly responded to a “catch” question within the questionnaires: “If you are paying attention to these questions, please select ‘A little’ as your answer” (N = 7). Combining all exclusion criteria, 42 participants (17.95%) were excluded. N = 192 participants were left for analysis (115 females (59.90%), between 18-65 ages (mean = 31.55, SD = 11.75 years).

### Two-step reinforcement learning task

We used the two-step reinforcement learning task (Daw et al., 2011, 2005) to assess individual differences in model-based planning. Participants had to navigate two stages to learn reward probabilities associated with options presented, with the main goal of earning rewards (‘space treasure’). The paradigm was presented with a cover story (***Figure 1***). In the first stage, participants had to choose between two spaceships, each with a higher probability (‘common’ transitions: 70%) of leading to one of two planets (second stage states, represented by coloured blocks) but that sometimes lead to the alternative (‘rare’ transitions: 30%) planet. Once on the planet, the participants then had to choose between two aliens which were probabilistically rewarded with either space treasure or no reward, indicated by a puff of ‘space dust’. Each alien, a total of four over two planets, had a unique probability of receiving space treasure, which drifted slowly and independently over time (always >0.25 or <0.75). Individuals performing in a model-based way would make decisions based on the history of rewards and their knowledge of the transition structure of the task, while individuals performing basic temporal difference (‘model-free’) learning would simply make decisions solely on the history of rewards obtained.

The sequence of events was presented in the same manner as a prior study that conducted the two-step task in the EEG (Eppinger et al., 2017) with the exception that we used the standard 70/30% transition probabilities (whereas Eppinger et al., 2017 instead contrasted blocks of 60/40% vs 80/20%) and had a slightly shorted time to make a choice (1500ms here versus 2000ms in their paper) (***Figure 1***). On each trial, participants were first presented with a fixation cross for 500ms, and then shown a choice between two spaceships. They had 1500ms to respond; after which, an outline over the chosen option would indicate their choice (feedback) for 500ms. A fixation cross was shown for 500ms before transition, where the transitioned planet was shown (a blank colour block) for 1000ms. Two aliens of that particular planet would then appear, with 1500ms for choice, with feedback of the chosen option subsequently shown for 500ms. Each of the aliens led to a probabilistic reward with a picture of ‘space treasure’, or no reward with ‘space dust’, that was presented for 1000ms. Responses were indicated using the left (‘Q’) and right (‘P’) keys. Colour of blocks behind rockets and those representing planets were randomised across all participants. Participants performed two blocks of 75 trials, i.e. 150 trials in total. Prior to the experimental task, participants completed a tutorial that explained the key concepts of the paradigm; the probabilistic association between the aliens and rewards (10 trials) and the probabilistic transition structure of rockets to planets (10 trials). After this practice phase, they had to answer a 3-item basic comprehension test regarding the key rules of the task. If participants failed to answer all questions correctly, the experimenter would reiterate the key concepts of the paradigm to the participant, allowing clarification.

### Behavioural data pre-processing

Individual missed trials and trials with very fast (<150ms) reaction times at the first stage (indicating inattention or poor responding) were excluded from analyses. A total of 1082 trials (3.76%) were removed across participants (per participant mean = 5.64 (3.76%) trials).

### Self-report psychiatric questionnaires, transdiagnostic dimensions & IQ

In order to quantify compulsivity in our sample, we applied a previously defined transdiagnostic definition (Gillan et al., 2016) that is based on a weighted combination of items drawn from 9 self-report questionnaires (which were fully randomized) assessing: *alcohol addiction* using the Alcohol Use Disorder Identification Test (AUDIT) (Saunders, Aasland, Babor, De La Fuente, & Grant, 1993), *apathy* using the Apathy Evaluation Scale (AES) (Marin, Biedrzycki, & Firinciogullari, 1991), *depression* using the Self-Rating Depression Scale (SDS) (Zung, 1965), *eating disorders* using the Eating Attitudes Test (EAT-26) (Garner, Bohr, & Garfinkel, 1982), *impulsivity* using the Barratt Impulsivity Scale (BIS-11) (Patton, Stanford, & Barratt, 1995), *obsessive-compulsive disorder* (OCD) using the Obsessive-Compulsive Inventory - Revised (OCI-R) (Foa et al., 2002), *schizotypy* scores using the Short Scales for Measuring Schizotypy (SSMS) (Mason, Linney, & Claridge, 2005), *social anxiety* using the Liebowitz Social Anxiety Scale (LSAS) (Liebowitz, 1987) and *trait anxiety* using the trait portion of the State-Trait Anxiety Inventory (STAI) (Spielberger, Gorsuch, Lushene, Vagg, & Jacobs, 1983). Correlations between questionnaire total scores ranged greatly (*r* = −0.08 to 0.79). A proxy for IQ was also collected using the International Cognitive Ability Resource (I-CAR) (Condon & Revelle, 2014) sample test which included 4 item types of three-dimensional rotation, letter and number series, matrix reasoning and verbal reasoning (16 items total). See ***Supplemental Figure S1*** for details of the distribution spread of the age, gender, IQ and questionnaire total scores.

We used weights derived from a previous study (Gillan et al., 2016) to transform the raw scores of the 209 individual items from the 9 questionnaires into dimension scores (‘Anxious-Depression’ (AD), ‘Compulsive Behaviour and Intrusive Thought’ (CIT; ‘compulsivity’), and ‘Social Withdrawal’ (SW)). This was because our sample size had too low a subject-to-variable ratio (N = 192) for *de novo* factor analysis, as compared to the original study (N = 1413). Prior studies have demonstrated the stability of the factor structure in new data (Rouault et al., 2018; Seow & Gillan, 2020). Consistent with prior work, the resulting dimension scores were moderately intercorrelated (*r* = 0.33 to 0.42).

### Quantifying model-based learning

Model-based estimates were estimated using mixed-effects models written in R, version 3.6.0 via RStudio version 1.2.1335 (http://cran.us.r-project.org) with the *glmer()* function from the *lme4* package, with Bound Optimization by Quadratic Approximation (bobyqa) with 1e5 functional evaluations. The basic model tested if participants’ choice behaviour to *Stay* (repeat a choice they made on the last trial) (stay: 1, switch: 0) was influenced by the previous trial’s *Reward* (rewarded: 1, unrewarded: −1), *Transition* (common (70%): 1, rare (30%): −1) and their interaction (***Supplemental Figure S2***). Within-subject factors (the intercept, main effects of reward, transition, and their interaction) were taken as random effects (i.e. allowed to vary across participants). In R syntax, the model was: Stay ~ Reward * Transition + (Reward * Transition + 1 | Subject). The extent to which model-based planning contributed to choice was indicated by the presence of a significant interaction effect between Reward and Transition (MB). Split half-reliability, where the data were split into two subsets (even versus odd trials) and correlated and adjusted with Spearman-Brown prediction formula, was estimated for model-based planning.

To test if the compulsive dimension was associated with model-based deficits, we included the total scores of all three dimensions (*AD*: anxious-depression, *CIT*: compulsive behaviour and intrusive thought, *SW*: social withdrawal) as z-scored fixed effect predictors into the basic model described above. The extent to which compulsivity is related to deficits in model-based planning was indicated by the presence of a significant negative Reward*Transition\**CIT* interaction. Prior work has shown that age and IQ are associated with model-based learning and to some extent compulsivity (Gillan et al., 2016); control analyses presented in ***Supplementary Materials*** demonstrate that these variables did not drive any of the results presented.

### Sensitivity to task structure: Reaction time (RT)

Recent work has shown that one effective way to index an individual’s sensitivity to the structure of the task is via reaction times (RT) (Shahar et al., 2019). In a similar fashion, we conducted a mixed effect linear regression of transition type (*Transition* (common: −1, rare: 1) on second stage reaction time (S2-RT). In the syntax of R, the model was: S2-RT ~ Transition + (Transition + 1 | Subject). We asked if compulsivity was associated with a reduction in RT sensitivity to the transition structure by including the total scores of the three dimensions (*AD, CIT, SW*) as z-scored fixed effect predictors into the original model above.

### EEG recording & pre-processing

EEG was recorded continuously using an ActiveTwo system (BioSemi, The Netherlands) from 128 scalp electrodes and digitized at 512 Hz. The data were processed offline using EEGLab (Delorme & Makeig, 2004) version 14.1.2 in MATLAB R2018a (The MathWorks, Natick, MA). Data were imported using A1 as a reference electrode, then downsampled to 250 Hz and band-pass filtered between 0.05 and 45 Hz. Bad channels were rejected with a criterion of 80% minimum channel correlation. All removed channels were interpolated, and the data were re-referenced to the average. To remove ocular and other non-EEG artefacts, ICA was run on continuous data with runica, *pca* option on, and its components were rejected automatically with Multiple Artifact Rejection Algorithm (MARA) (Winkler, Haufe, & Tangermann, 2011), an EEGLab toolbox plug-in, at a conservative criterion of >90% artefact probability. For all EEG analyses, other non-specific artefacts were removed after epoching using a criterion of any relevant electrode examined showing a voltage value exceeding ±100μV. If participants had a rate of >70% of total epochs failing this criterion, their data were excluded from all analyses (N = 4 as reported in **Participant exclusion criteria**). The remaining participants had mean = 147.46 (SD = 2.98) epochs left.

### Single-trial analyses with EEG signals

All analyses described below (including time-frequency single-trial analyses) were conducted with mixed effects models. For every single-trial analysis, we excluded single-trial EEG estimates which were within ±5 SD away from the mean of the group. A maximum of <0.79% (n = 215) of the total trials across all participants were excluded for any measure. The regression model-based estimate (MB) was used as the individual between-subjects model-based estimate in all EEG analyses.

### Sensitivity to task structure: P300 and transition type

We first measured the P300 component at four parietal electrodes over the topography of the stimulus-locked peak (D16 (CP1), A3 (CPz), B2 (CP2), A4); ***Figure 3***). Data were epoched from −500ms to 1700ms relative to the onset of the second stage stimulus (aliens presented) and baselined corrected from −200ms to 0ms. Stimulus-locked single-trial P300 amplitudes were estimated as the mean of ±100ms around the individual’s averaged latency of their positive peak within a search window 250ms to 1000ms after stimulus onset. To eliminate amplitude biases owing to latency variances due to RT, we subsequently aligned the epochs (measured at A4, A5, A19 (Pz), A32, the response-locked peak; ***Supplemental Figure S3***) to the time of choice response execution. The response-locked P300 amplitude was quantified as the mean amplitude −100ms to 0ms before response. We also measured the build-up rate of the response-locked signal as the slope of a straight line fitted to each single-trial waveform using the interval −400ms to −200ms. To investigate if the P300 was sensitive to rare versus common transitions and whether this depended on model-based control/compulsivity, we regressed both stimulus- and response-locked P300 measures against transition type *Transition*: rare: 1, common: 0) interacting with z-scored model-based estimates (*MB*) or compulsivity (*CIT*, controlled for the other psychiatric dimensions *AD* and *SW*), taking *Transition* and the intercept as random effects. In R syntax, the models were EEG ~ Transition*MB + (Transition + 1 | Subj) and EEG ~ Transition*(CIT + AD + SW) + (Transition + 1| Subj) respectively.

### Time-frequency analysis

EEG data were epoched for both first and second stages of the task for time-frequency analyses (alpha (9-13Hz) and theta (4-8Hz) power) detailed in the subsequent sections: −1700ms to 2200ms stimulus-locked at the first stage (rockets) as well as −2000ms to 3500ms stimulus-locked at second stage (aliens). Time-frequency calculations were computed using custom-written MATLAB (The MathWorks, Natick, MA) routines. The EEG time series in each epoch was convolved with a set of complex Morlet wavelets, defined as a Gaussian-windowed complex sine wave: *e^(−i2*time*f)^ e^(−time^2/2σ^2)^* where *i* is the complex operator, *time* is time, *f* is frequency, which increased from 2 to 40 Hz in 40 logarithmically spaced steps. σ defines the cycle (or width) of each frequency band and was set to cycle/2πf, where cycle increased from 4 to 12 in 40 logarithmically spaced steps in accordance with each increase in frequency step. The variable number of cycles leverages the temporal precision at lower frequencies and increases frequency precision at higher frequencies. From the resulting complex signals of every epoch, we extracted estimates of power. Power is defined as the modulus of the resulting complex signal: Ζ(*time*) (power time series: *ρ*(*time*) = real[*z*(*time*)]^2^ + imag[*z*(*time*)]^2^).

Stimulus-locked first stage epoch was baselined corrected to the average frequency power for each frequency band examined (i.e. alpha or theta) from −400ms to −100ms (corresponding to first stage fixation) while for stimulus-locked second stage epoch used −1400ms to −1100ms (corresponding to second stage fixation, before presentation of the coloured squares (planets)) as the baseline. The latter baseline window was chosen as the colour of the planets were predictive of the aliens; as such, choice-relevant neural activity may potentially merge in the interval between the onset of the planets and aliens. For single-trial estimates of frequency power, as baselining with division induces spurious power fluctuations due to trial-to-trial fluctuations, power at each individual trial was baselined corrected with the linear subtraction method with its corresponding baseline activity: (power(*time*) – power(*baseline*)), at each frequency, at each channel. For visualisation purposes in the figures presented, power was normalized by conversion to a decibel (dB) scale: (10*log10[power(*time*)/power(*baseline*)]).

### Sensitivity to task structure: Alpha power and transition type

Alpha power was measured at five parietal-occipital electrodes (A18, A19 (Pz), A20, A21, A31; surrounding A20 electrode; ***Supplemental Figure S4***) in an epoch centred on the onset of the second stage stimuli (aliens) (see **Time-frequency analysis**). Single-trial stimulus-locked alpha power estimates were measured as the mean power ±250ms around the average latency of the negative peak, specific for each individual, found within a search window 0ms to 1000ms after stimulus onset. We additionally obtained alpha power estimates quantified across four 1000ms rolling time bins by the mean amplitude within each time window. We labelled the time bins as they began from transition to the stimulus (alien presentation) from 0ms to 1000ms, followed by three windows spanning choice to reward from 1000ms to 2000ms, 2000ms to 3000ms, and 3000ms to 4000ms. The same approach of mixed effect models with **P300 and transition type** was used to examine the influence of model-based estimates/compulsivity on alpha power representation of rare versus common transitions, except for where *Transition* was coded differently (rare: −1, common: 1) for ease of interpreting the direction of interaction effects.

### Cognitive control: theta power during choice

For theta power (4-8Hz), power estimates were measured at four frontal midline electrodes (C21 (Fz), C22, C23 (FCz), A1 (Cz); see ***Supplemental Figure S6***) at the first stage (see **Time-frequency analysis**). The mean power ±250ms around the individual’s average latency of the positive peak found within a search window 0ms to 500ms after stimulus onset was taken for every epoch. We tested if single-trial theta power was associated with model-based estimates (*MB*) or to compulsivity (*CIT*, controlled for *AD* and *SW*) by taking them as z-scored main regressors against theta power.

### Specificity with psychiatric questionnaire scores versus transdiagnostic dimensions

Additionally, we examined the advantages of utilising a transdiagnostic definition of compulsivity as opposed to investigating single psychiatric questionnaires. We repeated the above time-frequency analyses (alpha and theta) with the individual total questionnaire scores (*QuestionnaireScore*, z-scored) replacing the three psychiatric dimensions (*CIT, AD, SW*) in their respective regression models detailed above. Separate mixed effects regression models were performed for each individual questionnaire as correlation across questionnaire scores ranged greatly from *r* = −0.09 to 0.79 as opposed to the transdiagnostic analysis where all three dimensions (that correlated moderately: *r* = 0.33 to 0.42) were included in the same model.

### Supplemental analyses

Finally, to ensure the specificity of any observed effects to the task events outlined above, we also tested for an association between model-based planning and compulsivity with our candidate EEG signatures in reverse. That is, we tested if between model-based planning and compulsivity were linked to (i) alpha power at the first stage or (ii) theta power sensitivity to transition type at the second stage. See ***Supplemental Figure S4*** and ***Supplemental Figure S6*** for the respective analyses.

## Code and data availability

The code and data to reproduce the main figures of the paper are freely available at https://osf.io/mx9kf/.

## Author Contributions

TXFS: Conceptualization, Methodology, Investigation, Formal analysis, Writing – Drafting, Review & Editing. RO: Methodology, Writing – Review & Editing. CMG: Conceptualization, Methodology, Writing – Drafting, Review & Editing, Supervision.

## Acknowledgments

This work is supported by a Postgraduate Ussher fellowship from Trinity College Dublin to TXFS and a ‘Institutional Strategic Support Fund’ grant (204814/Z/16/A) to Trinity College Dublin funded by the SFI-HRB-Wellcome Trust partnership. We would like to thank Edith Benoit, Caoimhe Dempsey, Maeve Jennings and Aoibheann Maxwell for assisting with data collection.

## Supplementary Materials

### Controlling for age and IQ

Age and IQ are known to covary with model-based learning (Gillan et al., 2016). Here, only IQ was significantly associated model-based learning (*β* = 0.12, *SE* = 0.03, *p* < 0.001); age was not (*β* = −0.04, *standard error (SE)* = 0.03, *p* = 0.30). Nonetheless, inclusion of both variables did not change the pattern of our main findings. Reduced goal-directed control was linked to compulsivity (*β* = −0.08, *SE* = 0.04, *p* = 0.03) and individuals high in model-based control showed larger alpha power differences between the two transition types over the three rolling time bins beginning from the transition (planet) (0ms to 1000ms: *β* = 0.03, *SE* = 0.02, *p* = 0.06) to the end of choice feedback (1000ms to 2000ms: *β* = 0.03, *SE* = 0.01, *p* = 0.04; 2000ms to 3000ms: *β* = 0.05, *SE* = 0.02, *p* = 0.05).

**Supplemental Table S1.**
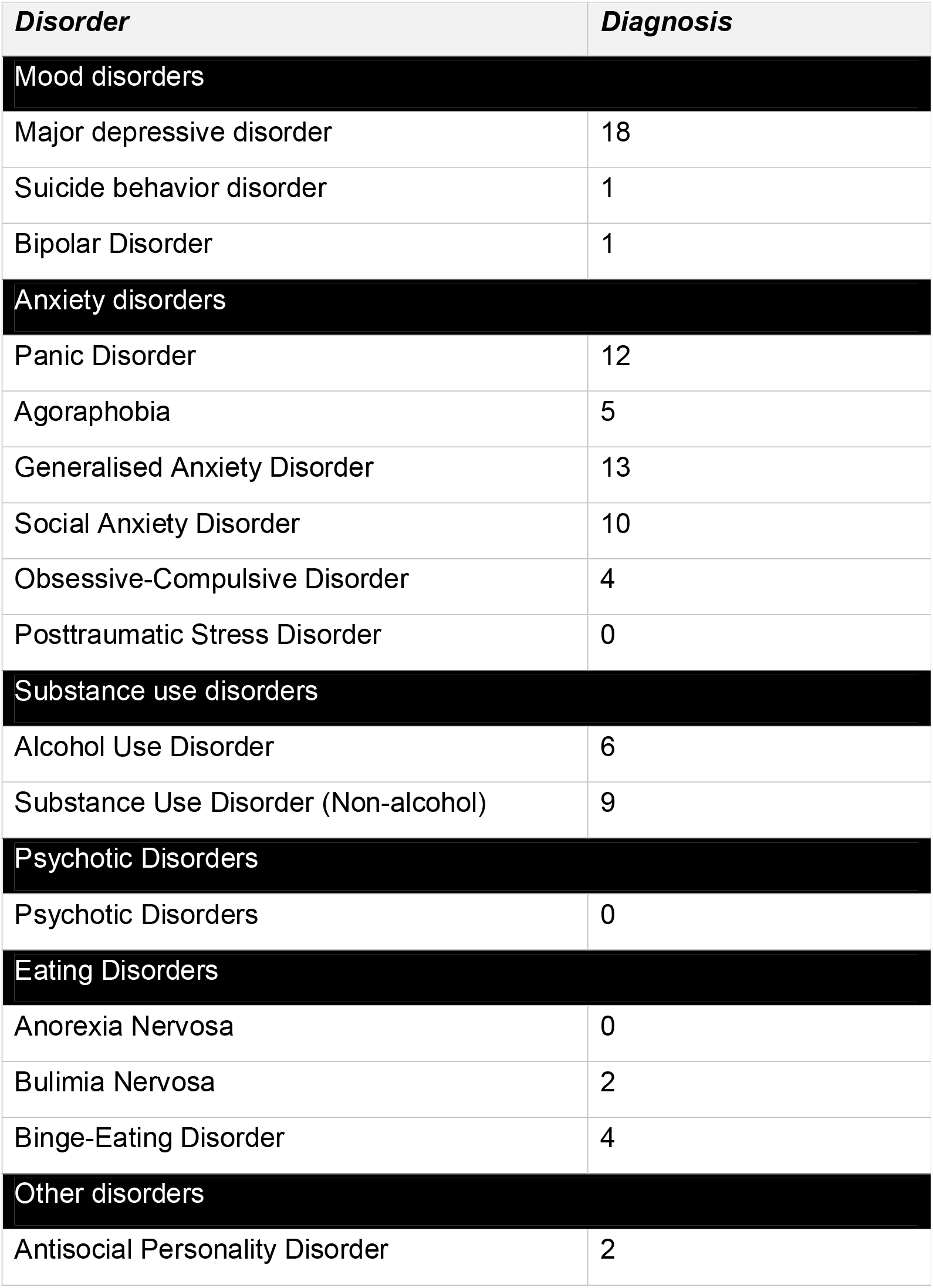
Mini International Neuropsychiatric interview (M.I.N.I.) diagnostic information summary for participants who presently met the criteria for at least one DSM-V disorder (N = 35).

### Disorder prevalence (M.I.N.I.)

After exclusion, 80 participants (41.67%) completed the M.I.N.I., which was introduced part-way through the study to add additional clinical context above our self-report measures. Of these participants, 35 (43.75%) met the criteria for one or more disorder. Broken down by recruitment arm, all 7 subjects (100%) recruited from the clinical setting met criteria, while 28 (38.36%) from university channels met criteria. This rate is close to published reports on the prevalence of mental health disorders in college student samples (Auerbach et al., 2018; Evans, Bira, Gastelum, Weiss, & Vanderford, 2018). Of the total sample, 33 (17.19%) were currently medicated for a mental health issue. Broken down by recruitment arm, all individuals recruited from the clinic were medicated, while 26 (14.05%) of those recruited through normal channels were medicated.

**Supplemental Figure S1.**
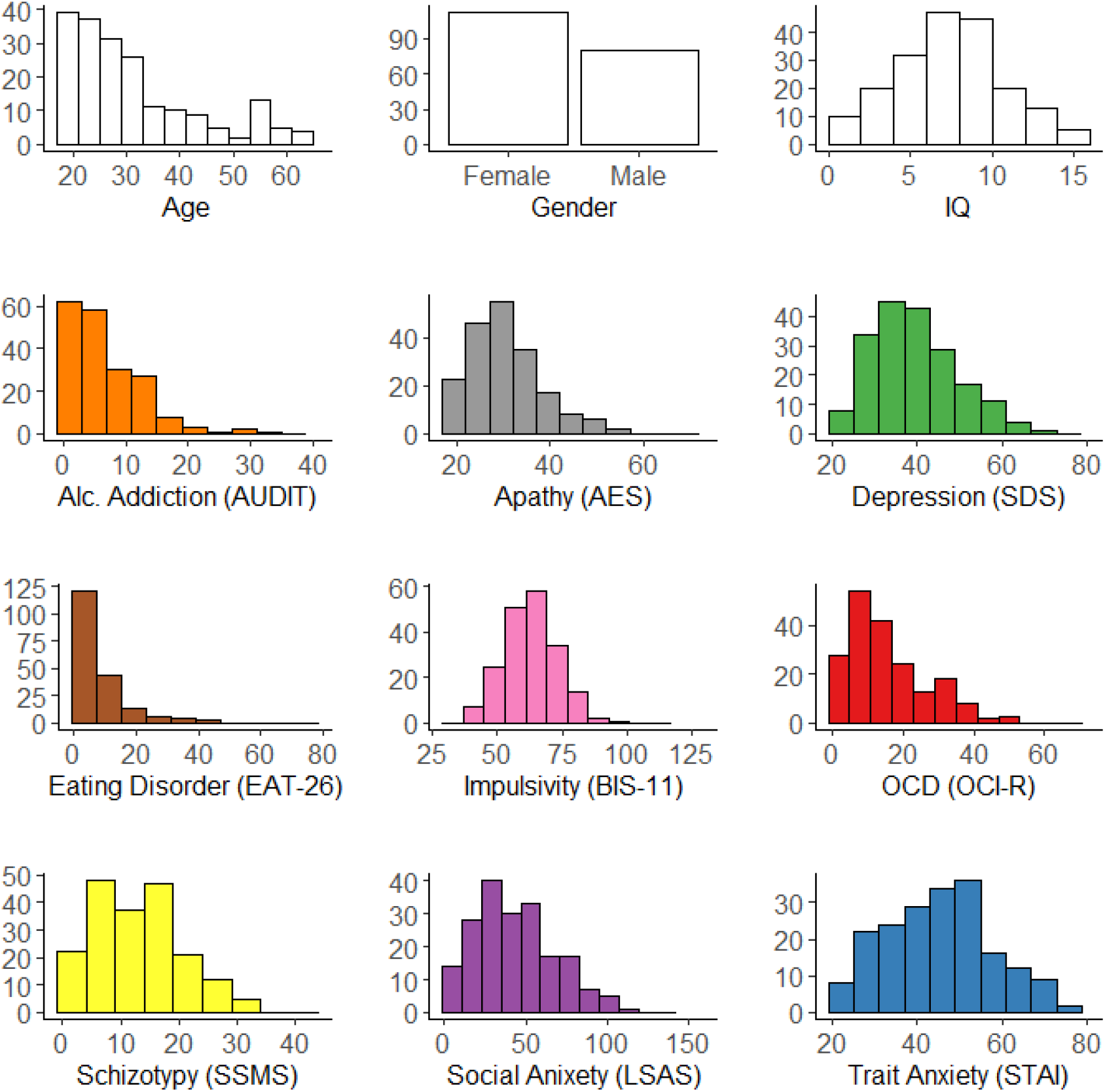
Histogram of demographics (age, gender and IQ) and total questionnaire scores across participants. Y-axes of each plot indicates the number of participants.

**Supplemental Figure S2.**
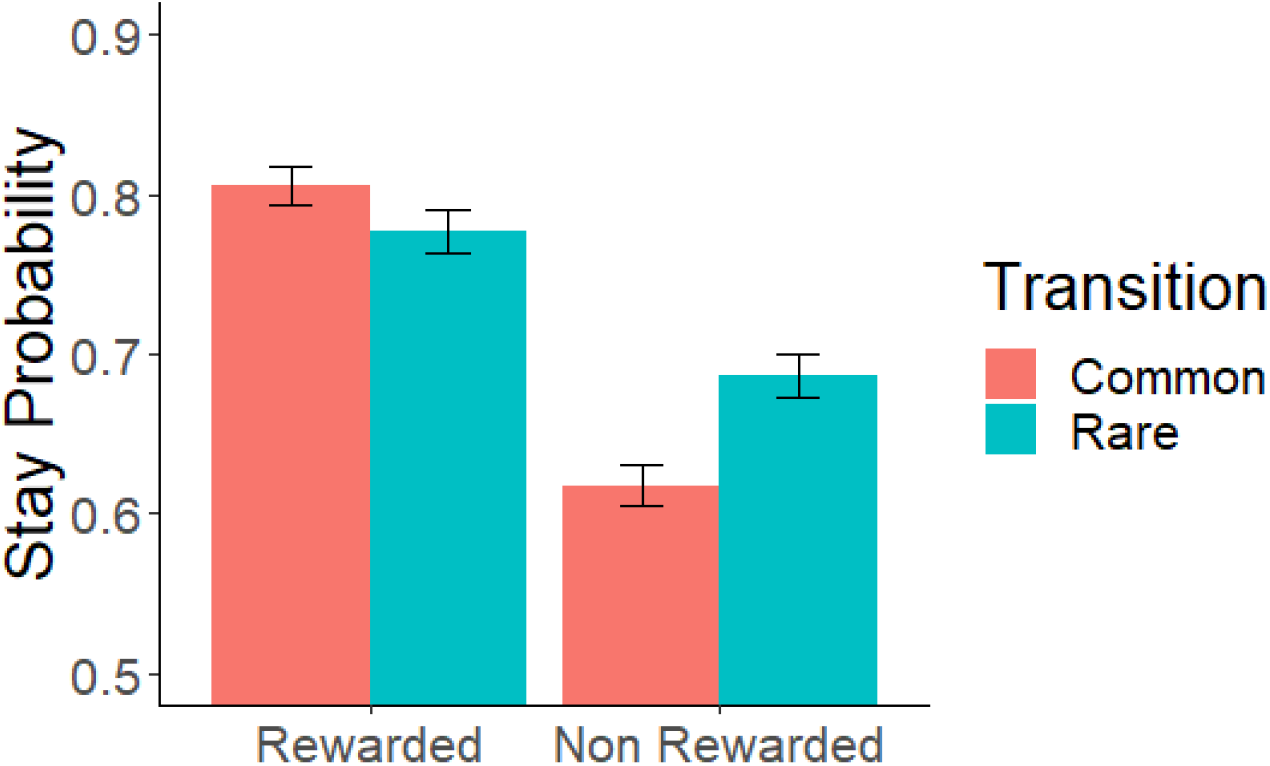
First stage stay probabilities. Model-based behaviour is reflected as the probability of repeating the first stage choice (stay) as a function of the occurrence of a transition from the previous trial (common: 70%, rare: 30%) and whether a reward was received (reward, non reward). In a purely model-free learner, stay probabilities after reward should be higher than when no reward was presented regardless of transition type. In a purely model-based learner, stay probabilities after common-reward and rare-non reward should be higher than common-non reward and rare-reward. In our empirical data here, the stay probabilities obtained across conditions is a mix of both model-based and model-free behaviour. Error bars reflect standard errors of mean.

**Supplemental Figure S3.**
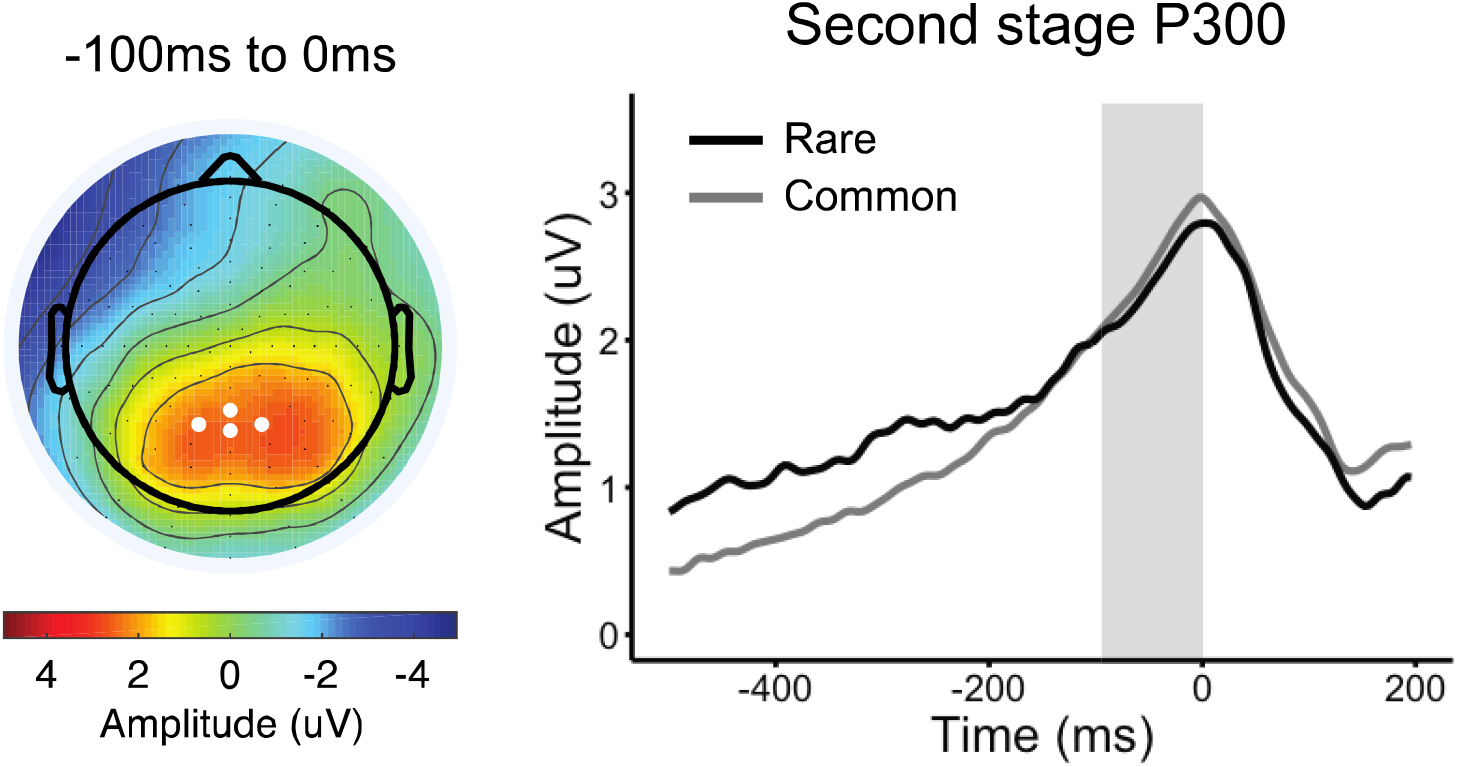
Response-locked P300 and transition type. Topography plot represents the P300 component −100ms to 0ms before second stage response. White dots indicate parietal electrode sites (A4, A5, A19 (Pz), A32) where the positive component was measured. Grand average second stage P300 is plotted response-locked comparing the waveforms following rare versus common transitions. Single-trial analyses indicate that the P300 amplitude, measured as the mean amplitude −100ms to 0ms (shaded grey), does not distinguish transition type (β = −0.09, SE = 0.08, p = 0.23).

**Supplemental Figure S4.**
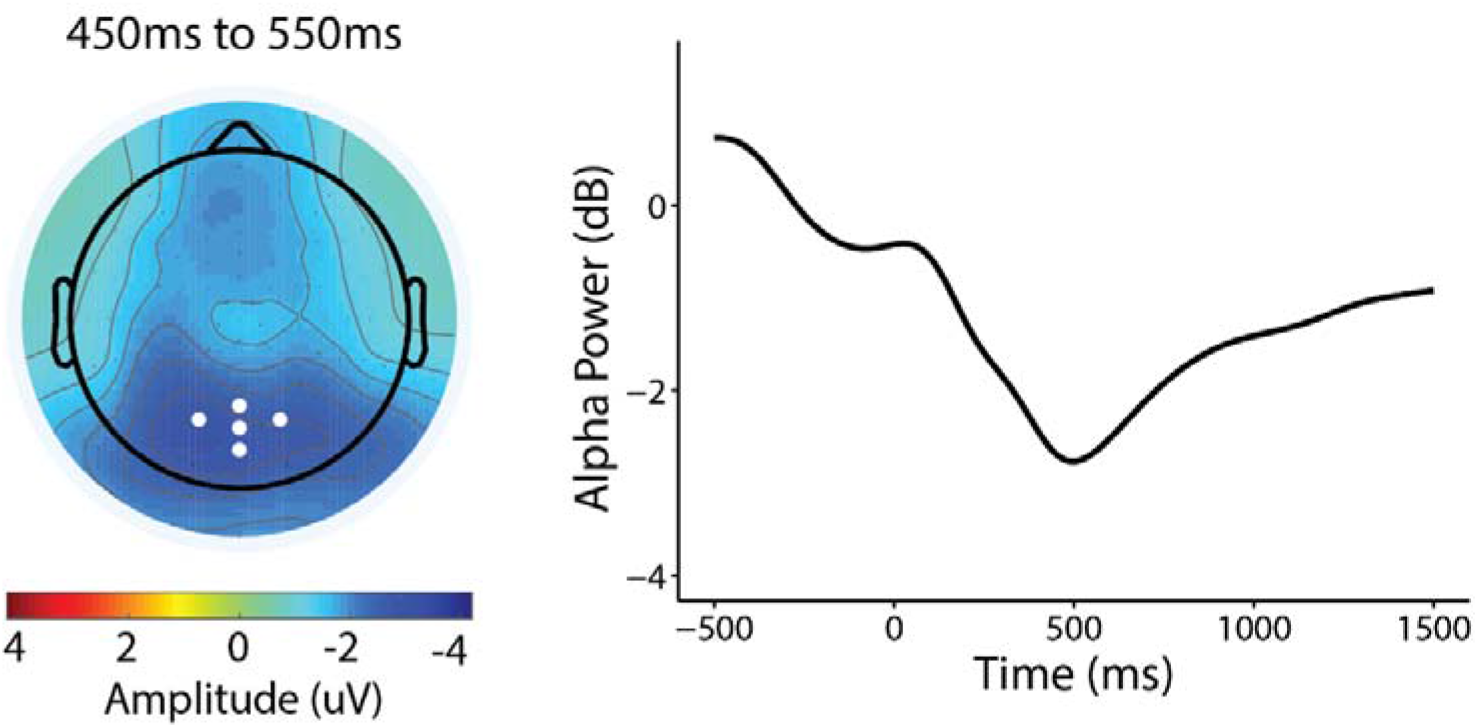
First stage alpha power. Topography and line plot (locked to first-stage rockets) show alpha depression during the making a choice at the first stage. White dots on the topography plot indicate parietal-occipital electrode sites (A18, A19 (Pz), A20, A21, A31) where alpha was measured for both first and second stages.

**Supplemental Figure S5.**
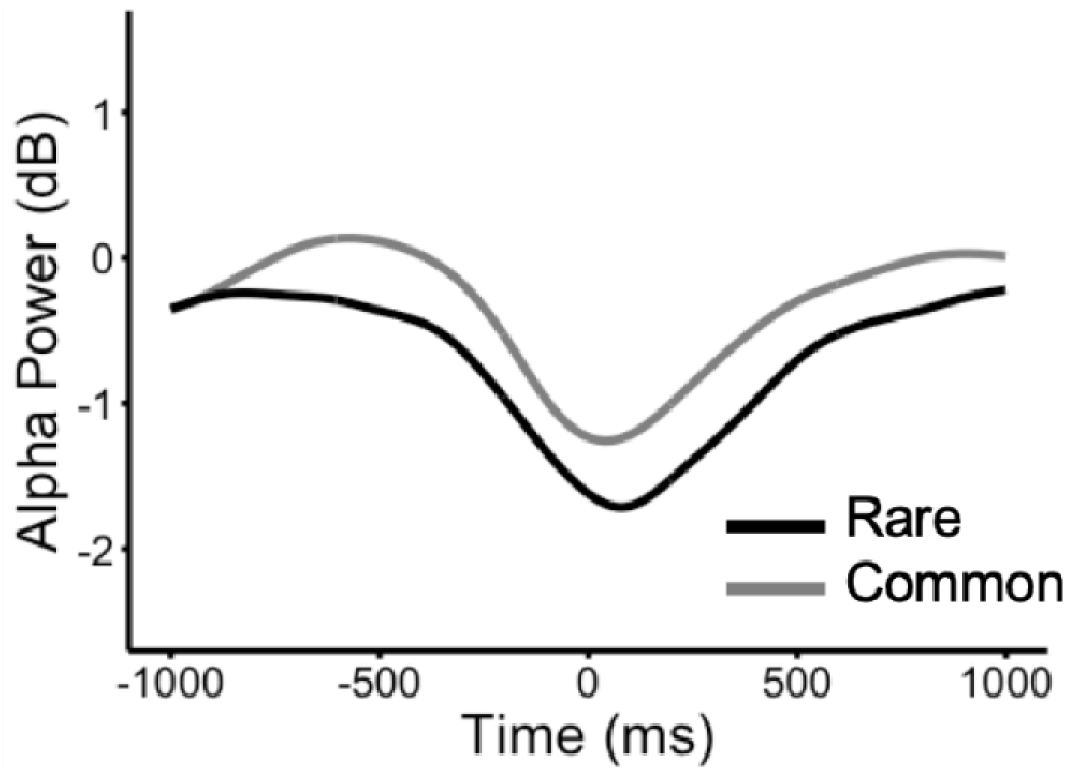
Grand average waveforms of rare versus common transitions for second stage response-locked alpha power.

### Alpha transition effect is not explained by RT

RT differences between rare and common transitions was only significantly associated with stimulus-locked alpha power differentiation of states in the time bin before reward presentation (2000ms to 3000ms: *β* = 0.03, *SE* = 0.02, *p* = 0.04; all other time bins: *p*s > 0.30; ***Figure 4***). To complement our main result based on stimulus-locked alpha, we repeated the transition analysis with single-trial response-locked alpha estimates (measured as the mean of ±100ms centered around each participant’s averaged latency of the negative peak), which also yielded a significant association overall effect (*β* = 0.06, *SE* = 0.01, *p* < 0.001; ***Supplemental Figure S5***). Similar to stimulus-locked alpha, rare transitions showed greater depression of alpha during choice selection for rare versus common transitions.

**Supplemental Figure S6.**
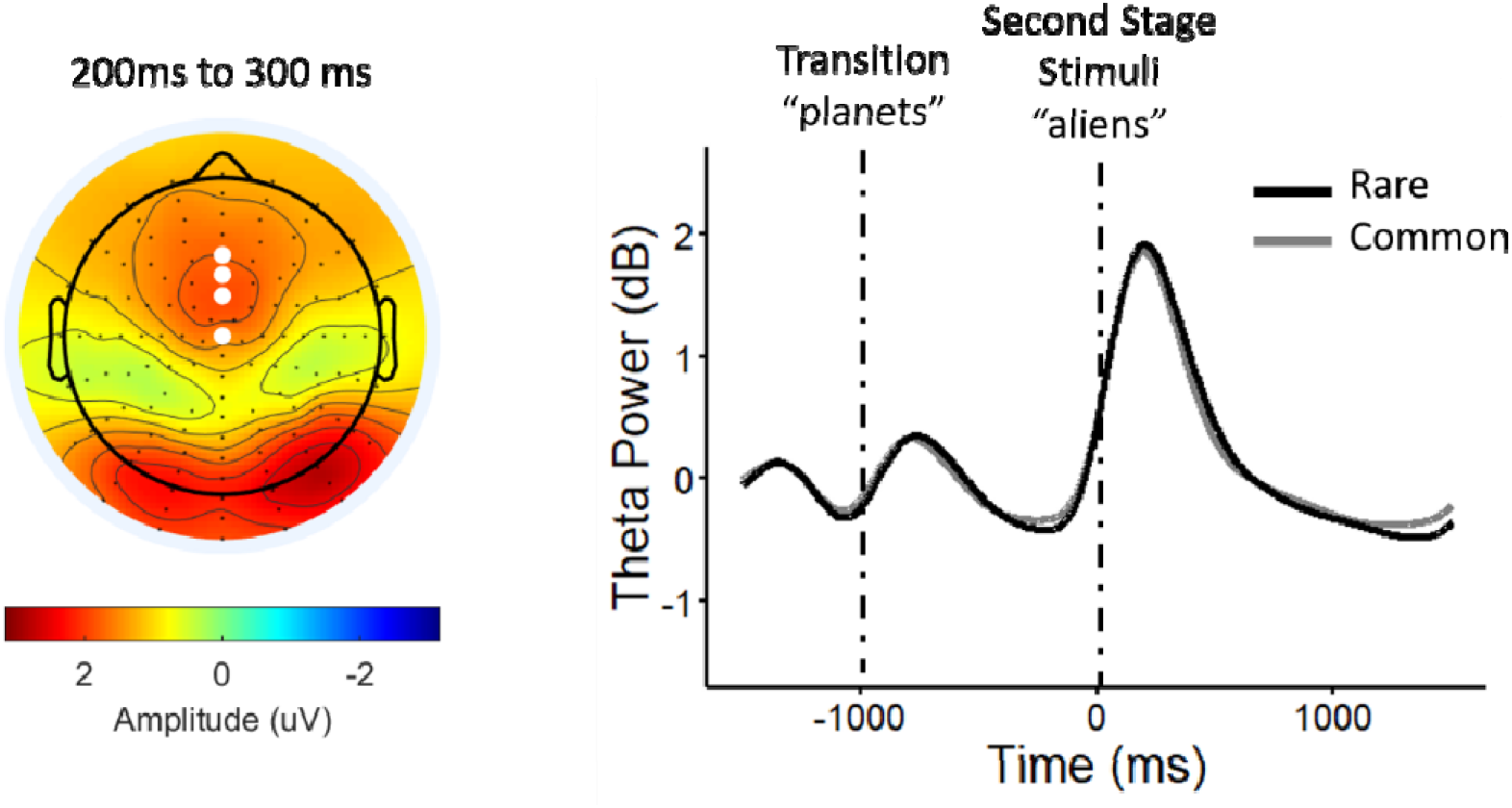
Second stage stimulus-locked theta power. Topography plot shows theta power increase after stimulus-onset at the mid-frontal scalp. White dots indicate electrode sites (C21 (Fz), C22, C23 (FCz), A1 (Cz)) where theta power was measured. Theta power was not associated to compulsivity (β = −0.004, SE = 0.02, p = 0.84) nor model-based planning (β = 0.01, SE = 0.02, p = 0.51). Theta power was also not linked to transition type (β = −0.01, SE = 0.01, p = 0.20) and had no transition interaction effects with compulsivity (β = 0.02, SE = 0.03, p = 0.14) nor model-based planning (β = −0.004, SE = 0.01, p = 0.65).

